# Anti-inflammatory treatment with FTY720 starting after onset of symptoms reverses synaptic and memory deficits in an AD mouse model

**DOI:** 10.1101/2019.12.15.868026

**Authors:** Georgia-Ioanna Kartalou, Ana Rita Salgueiro Pereira, Thomas Endres, Angelina Lesnikova, Plinio Casarotto, Paula Pousinha, Kevin Delanoe, Elke Edelmann, Eero Castrén, Kurt Gottmann, Helene Marie, Volkmar Lessmann

**Affiliations:** Institute of Physiology, Medical Faculty, Otto-von-Guericke University, 39120 Magdeburg, Germany; Institute of Neuro- and Sensory Physiology, Medical Faculty, Heinrich Heine University Duesseldorf, Germany; Université Côte d’Azur, CNRS, IPMC, UMR7275 Valbonne, France; Neuroscience Center, HiLIFE, University of Helsinki, Helsinki, Finland; Center for Behavioral Brain Sciences (CBBS), 39120 Magdeburg, Germany

**Keywords:** Alzheimer’s disease, fingolimod, FTY720, APP/PS1, spines, LTP, spatial memory, microglia, astrogliosis, neuro-inflammation, BDNF, hippocampus, learning and memory, neurodegenerative disease

## Abstract

Therapeutical approaches providing effective medication for Alzheimer’s disease (AD) patients after disease onset are urgently needed. Repurposing FDA approved drugs like fingolimod (FTY720) for treatment of AD is a promising way to reduce the time to bring such medication into clinical practice. Previous studies in AD mouse models suggested that physical exercise or changed lifestyle can delay AD related synaptic and memory dysfunctions when treatment started in juvenile animals long before onset of disease symptoms. Here, we addressed whether the FDA approved drug fingolimod rescues AD related synaptic deficits and memory dysfunction in an APP/PS1 AD mouse model when medication starts after onset of symptoms (at 5 months). Male mice received intraperitoneal injections of fingolimod for 1-2 months starting at 5-6 months. This treatment rescued spine density as well as long-term potentiation in hippocampal CA1 pyramidal neurons, and ameliorated dysfunctional hippocampus-dependent memory that was observed in untreated APP/PS1 animals at 6-7 months of age. Immunohistochemical analysis with markers of microgliosis (Iba1) and astrogliosis (GFAP) revealed that our fingolimod treatment regime strongly down regulated neuro-inflammation in the hippocampus and cortex of this AD model. These effects were accompanied by a moderate reduction of Aβ accumulation in hippocampus and cortex. Our results suggest that fingolimod, when applied after onset of disease symptoms in an APP/PS1 mouse model, rescues synaptic pathology and related memory performance deficits observed in untreated AD mice, and that this beneficial effect is mediated via anti-neuroinflammatory actions of the drug on microglia and astrocytes.

## Introduction

Alzheimer’s disease (AD) is the most common form of dementia, characterized by a progressive decline of cognitive functions (Blennow et al., 2006). Recent research has seen significant advances in better understanding the cellular and molecular mechanisms of AD. This includes the general consensus that Aβ pathology that starts in the human neocortex 10-15 years before AD becomes symptomatic, and hippocampal tau pathology (which coincides with early stages of symptomatic AD in humans), together, underlie AD disease onset (Livingston et al., 2017). One of the main objectives of AD research is to identify novel therapeutic approaches alleviating cognitive deficits. While life style factors like e.g. physical exercise and a healthy diet might be able to delay the onset of AD, no treatment for AD has thus far been reported that can effectively counteract dementia symptoms after disease onset. Importantly, neuroinflammatory signals mediated by microglial cells and concomitant astrogliosis are major hallmarks of AD (Heneka et al., 2015; Sarlus and Heneka, 2017). Accordingly, therapeutic approaches focusing on reduction in neuroinflammatory signaling of microglial cells are among the most promising intervention strategies to ameliorate AD deficits. Therefore, repurposing anti-inflammatory drugs for AD therapy that are already approved by the FDA for treatment of human subjects could provide an important benefit to rapidly develop an effective AD medication.

Fingolimod (FTY720) is an FDA approved drug for the treatment of multiple sclerosis (MS) in humans since 2011 and has evolved in recent years as the first line oral medication for relapsing MS in clinical practice (Druart et al., 2018). In target tissue, it can be converted to fingolimod phosphate, which is a potent modulator of sphingosine-1-phosphate receptors (S1PR_1-5_; (Angelopoulou and Piperi, 2019; Chaudhry et al., 2017). In the immune system, fingolimod induced S1PR receptor modulation promotes the retention of T-lymphocytes within lymph nodes, thereby reducing the invasion of the CNS by lymphocytes where they promote inflammatory processes leading to demyelination (Chaudhry et al., 2017). More recently, fingolimod was shown to increase BDNF mRNA and protein levels in mouse models of different neurodegenerative diseases (Deogracias et al., 2012; Di Pardo et al., 2014; Fukumoto et al., 2014; Miguez et al., 2015). Deficits in BDNF signaling are discussed to contribute to cognitive dysfunctions in AD (Amoureux et al., 1997; Murray et al., 1994; Phillips et al., 1991), and enhanced signaling through the BDNF receptor TrkB can ameliorate cognitive symptoms in AD model mice (Kemppainen et al., 2012; Rantamaki et al., 2013). Given the anti-inflammatory effects of fingolimod and the potentially beneficial effects of this FDA approved drug on BDNF/TrkB signaling we set out to explore the capacity of fingolimod to be repurposed for the treatment of AD-like pathology in an APP/PS1 Alzheimer mouse model. Importantly, we started with a 1-2 months long fingolimod treatment period at 5-6 months of age, when these mice had already entered the symptomatic phase of the disease. Our results reveal rescue of spine deficits, as well as of impaired LTP and memory dysfunction in this AD mouse model. These effects occurred independent from BDNF signaling but are consistent with the anti-inflammatory actions of fingolimod on microglia and astrocytes that we observed. Hence, our results reveal a high potential of fingolimod as an efficient drug for treatment regimes starting after AD symptoms have been diagnosed.

## Results

We used a previously established double transgenic APP/PS1 mouse model that starts to develop amyloid-β (Aβ) pathology in the neocortex and hippocampus at 2 and 4 months of age, respectively (Radde et al., 2006). Impaired LTP at Schaffer collateral CA1 synapses in acute slices *ex vivo* is observed at 5 months in these mice (**Fig. 1**) and becomes detectable with *in vivo* recordings beyond 6 months of age (Gengler et al., 2010). Moreover, hippocampus-dependent learning deficits in these APP/PS1 mice become apparent 5 months after birth (**Fig. 1**; (Psotta et al., 2015; Radde et al., 2006)). To test for a possible amelioration of the AD phenotype at a symptomatic stage in our APP/PS1 mouse model, we started fingolimod (FTY720) treatment at 5-6 months of age, when AD-like symptoms are present. Male APP/PS1 mice and littermate controls were treated with intraperitoneal (i.p.) injections of fingolimod (FTY720; 1.0 mg/kg body weight) every second day. Spine density and Schaffer collateral LTP in CA1 hippocampal neurons, as well as hippocampus-dependent learning were tested at 6-7 months (i.e. after 4-8 weeks of fingolimod treatment).

**Fig. 1:**
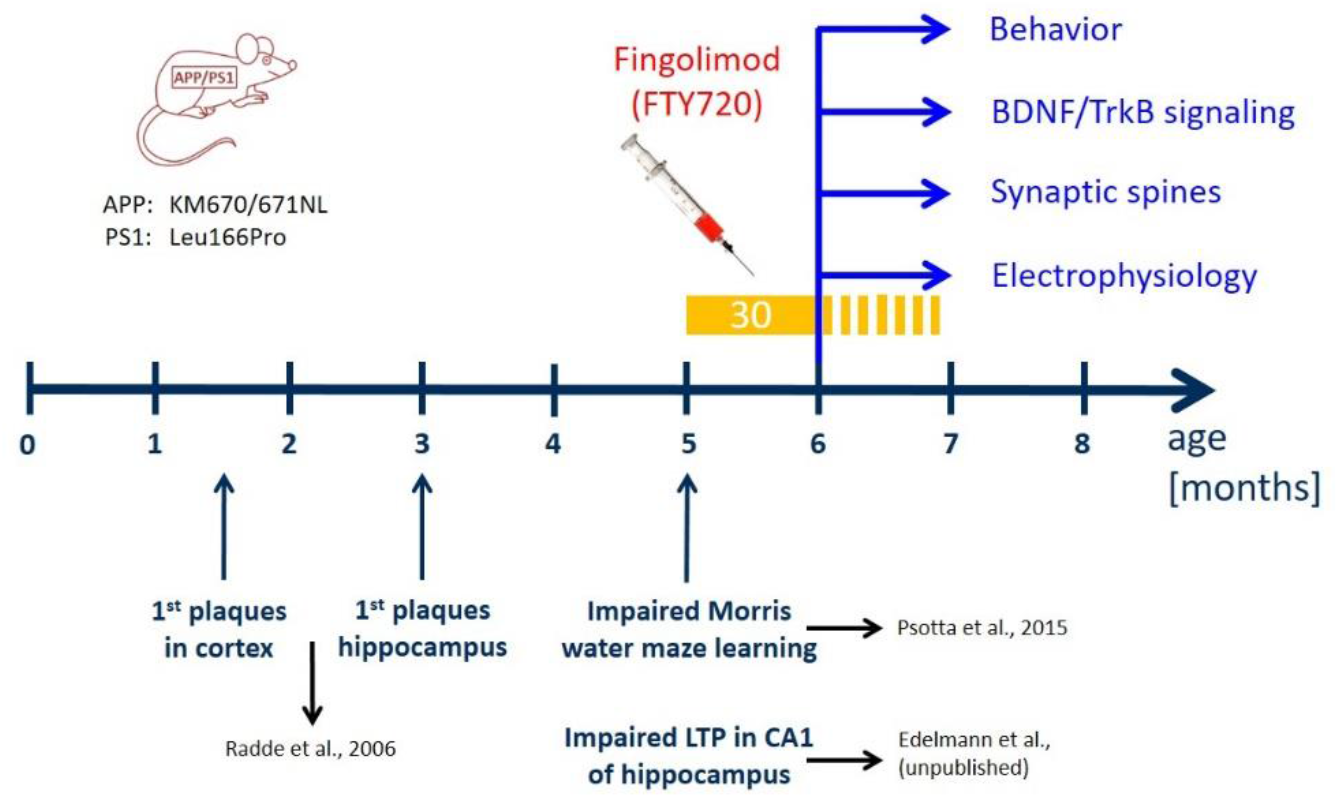
Time line of treatment with fingolimod (FTY720) compared to development of AD-like pathology in the APP/PS1 mouse model used in the study. Mutations in APP and presenilin 1 in the APP/PS1 mouse model used in our study are indicated on the left (Radde et al., 2006). In these APP/PS1 mice, Aβ plaques are detectable after 6 weeks in the cortex and after 3 months in the hippocampus (Radde et al., 2006). Moreover, these APP/PS1 mice show spatial learning deficits in the Morris water maze (Psotta et al., 2015) and impaired LTP at Schaffer collateral-CA1 synapses in the hippocampus (Edelmann et al., data not shown) at 5 months of age (i.e. starting at 22 weeks). FTY720 injections (every second day; yellow bar) started at 5 months (no earlier than 22 weeks after birth) when the animals already display synaptic and memory deficits. The injections were carried on for at least 30 days thereafter, and were continued (stippled yellow bar) until completion of behavioral testing, or until animals were sacrificed for analyses (spines, LTP, immunohistological stainings, biochemical BDNF and TrkB signaling analysis) at 6-7 months.

### Chronic fingolimod (FTY720) treatment of 5-6 months old APP/PS1 mice rescues spine deficits

Golgi-Cox staining was used to determine spine densities of CA1 pyramidal neurons in secondary apical dendrites located in stratum radiatum of the hippocampus. To directly relate spine numbers to Aβ pathology, we labelled Aβ plaques in 6-7 months old animals by i.p. injecting animals twice with the blue fluorescent dye Methoxy X-04 (Liebscher et al., 2014)(Kartalou et al., in prep.) one day before and 2 hours before sacrificing the mice for the Golgi staining procedure. In this set of experiments, dendritic stretches analyzed for spine density where subdivided into segments with a minimal distance of spines <50 μm (i.e., 3 - 44 μm) to the closest Aβ plaque (termed: near), and segments with 50-250 μm (i.e., 54 - 227 μm) minimal distance to Aβ plaques (termed: distant). Since plaque load is low in CA1 and stratum radiatum of the hippocampus at this AD stage (compare Fig. 6A,B), there was no second plaque closer than 300 μm to the analyzed dendrite. Spine analysis in these two groups revealed significantly reduced spine densities in APP/PS1 animals that was more pronounced when spines were located near to Aβ plaques (WT: 1.78 ± 0.02; APP/PS1 near: 1.31 ± 0.03; APP/PS1 distant: 1.46 ± 0.03; **Fig. 2**).

**Fig. 2:**
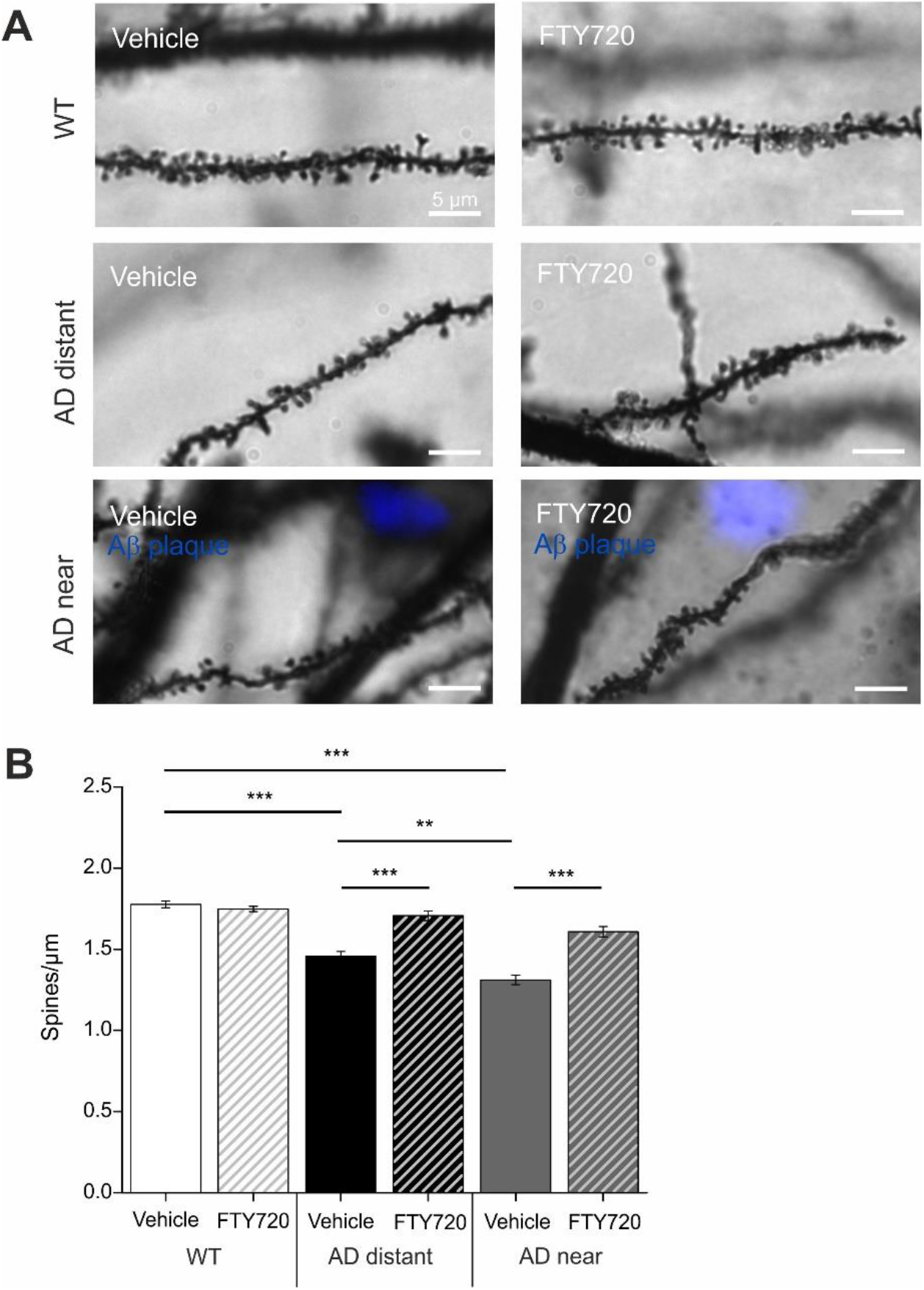
Chronic fingolimod (FTY720) treatment rescues spine deficits in APP/PS1 mice. 5-6 months old male APP/PS1 mice were treated with i.p. injections of fingolimod (FTY720) for 1-2 months. Spine density was determined with Golgi-Cox staining in hippocampal CA1 pyramidal neurons at 6-7 months. In APP/PS1 mice, Aβ plaques were visualized with blue fluorescent Methoxy X04. **(A)** Secondary apical dendrites of CA1 pyramidal neurons in vehicle and fingolimod treated WT animals (WT, upper panel), in APP/PS1 animals >50 μm away from the closest plaque (AD distant, middle panel), or in APP/PS1 mice <50 μm away from the nearest plaque (AD near, lower panel). **(B)** Quantification of spine densities in secondary apical CA1 dendrites shown for the 6 different groups in (A). Note the complete rescue of spine deficits in dendrites distant to plaques in fingolimod treated APP/PS1 mice, and the significant amelioration of spine deficits in dendrites close to plaques. Horizontal bars indicate statistical significance between selected groups. Significance of differences was tested with two-way ANOVA followed by Tukey’s post hoc test (n=30 dendrites from 3 animals per group). Significance level was set to 0.05 (p<0.05). Different levels of significance are indicated by stars, with ** = p< 0.01, ***= p<0.001.

Strikingly, 1-2 months treatment of mice with fingolimod (starting at 5-6 months of age) completely rescued spine densities of dendrites distant to plaques back to the level of WT littermate controls, and significantly ameliorated the spine deficits in dendrites near plaques (APP/PS1 + FTY720 near: 1.61 ± 0.03; APP/PS1 + FTY720 distant: 1.71 ± 0.03, post hoc Tukey’s test with p<0.0001). Overall, two-way ANOVA analysis revealed a significant main effect of genotype x treatment interaction (F (2, 174) = 21.51 P<0.0001). Of note, the identical fingolimod treatment of WT littermates did not change CA1 pyramidal neuron spine densities in these animals (WT vehicle: 1.78 ± 0.02; WT + FTY720: 1.75 ± 0.02).

These results demonstrate that chronic treatment of APP/PS1 mice with fingolimod starting with onset of disease symptoms at 5-6 months drastically reduced spine pathology in this Alzheimer mouse model.

### Chronic application of fingolimod (FTY720) rescues LTP deficits at CA3-CA1 synapses in APP/PS1 mice

Extracellular field potentials were recorded to analyze long-term potentiation (LTP) in acute hippocampal slices from 6-7 months old APP/PS1 mice and WT littermate controls. The recording electrode was placed in stratum radiatum of the CA1 area, and LTP was elicited by high frequency tetanic stimulation of Schaffer collateral axons projecting from the CA3 area. Sixty minutes after LTP induction, the percentage of potentiation was determined in the four groups. The two-way ANOVA showed a significant main effect of genotype x treatment interaction (F(1,31)=19.59 P=0.0001), but no main effects of genotype or treatment. Vehicle treated WT littermates showed robust potentiation of field EPSP slopes (WT vehicle: 133.4 ± 4.65%; **Fig. 3A, B**), whereas in hippocampal slices obtained from vehicle treated APP/PS1 mice LTP was significantly reduced (APP/PS1 vehicle: 107.6 ± 3.2%, n=8, significantly different from WT, Tukey comparison p<0.01; **Fig. 3A, B**). Importantly, chronic treatment of AD mice with FTY720 completely rescued LTP when compared to vehicle-treated AD mice (AD FTY720: 135.9 ± 6.1%, n=8, significantly different from AD vehicle, Tukey’s p<0.01). Fingolimod treatment of WT littermates reduced LTP magnitude, however, this alteration did not reach significance when compared to WT vehicle (WT + FTY720: 116.8 ± 5.7%, WT vehicle compared to WT FTY720 mice p>0.05). While the latter result suggests a trend of fingolimod towards decreasing LTP magnitude in WT animals, this does not harm the complete LTP rescuing effect of the same treatment in APP/PS1 mice. Since fingolimod treatment did in either genotype not affect the extent of the typical synaptic fatigue during high frequency synaptic stimulation used for LTP induction, and also did not affect the degree of post-tetanic potentiation (PTP; **Suppl. Fig. 1**), these data suggest that fingolimod affects LTP expression mechanisms rather than LTP induction processes in our recordings.

**Fig. 3:**
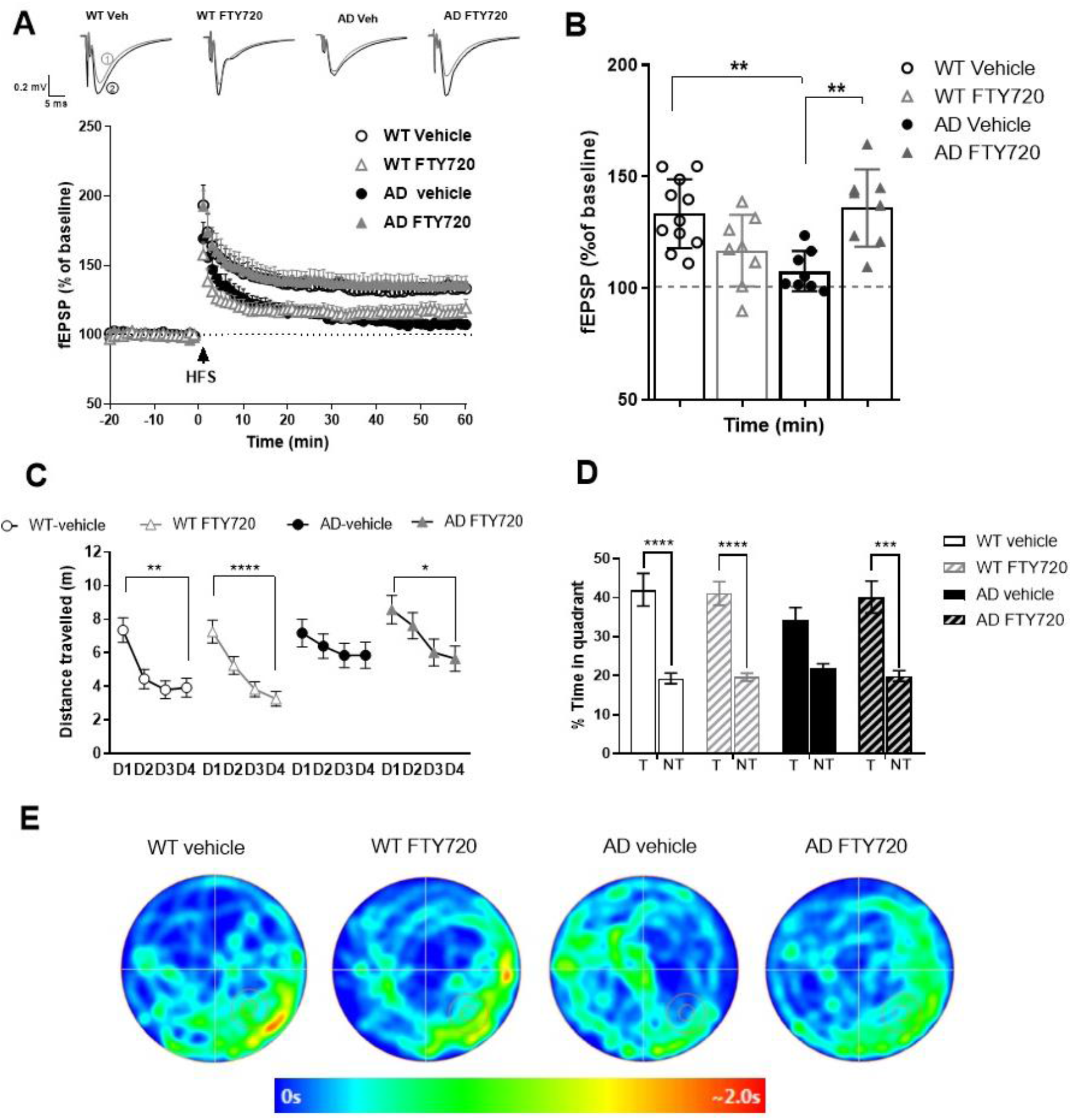
Chronic treatment with fingolimod (FTY720) rescues LTP and ameliorates spatial memory deficits in APP/PS1 mice. **(A, B)** Long-term potentiation evoked by high-frequency stimulation (HFS) in 6-7-month old WT and APP/PS1 mice after chronic vehicle or FTY720 treatment. **(A)** The graphs show the time-course of slope (normalized to baseline) of field excitatory post-synaptic potentials (fEPSPs) before and after induction of LTP with high frequency stimulation (HFS). Representative traces for each condition are shown before (1 - grey) and 60 min after LTP induction (2 - black). **(B)** Average of fEPSP slopes between 45-60 minutes after HFS. WT Vehicle (n=11), AD Vehicle (n=8), WT FTY720 (n=8) and AD FTY720 (n=8). **(C-E)** Spatial learning and memory retrieval after chronic vehicle or FTY720 treatment. **(C)** Spatial learning training in Morris Water Maze from training day 1 (D1) to training day 4 (D4) represented as distance travelled to the platform. **(D)** Memory recall (probe trial), 24 hours after the last training day (D4), represented as percent time spent in target (T) and average of non-target quadrants when platform was removed. WT vehicle (n=18), AD vehicle (n=15), WT FTY720 (n=20) and AD FTY720 (n=16). Horizontal bars indicate statistical significance between selected groups. **(E)** Heat maps represent the average occupancy time of different locations in the maze (red: high; blue: low residence time) for a set of animals from each group. Target quadrant (bottom right) is marked with the platform zone circle. Significance of differences was tested with two-way ANOVA. Different levels of significance are indicated by stars: * p<0.05, ** p<0.01, *** p<0.001; **** p<0.001.

Together, these electrophysiological data suggest that Schaffer collateral – CA1 LTP is completely rescued in APP/PS1 mice by the identical regime of fingolimod treatment that also rescues spine deficits.

### Chronic fingolimod (FTY720) treatment ameliorates memory deficits in 6-7 months old APP/PS1 mice

Hippocampus-dependent memory was tested in a Morris water maze spatial memory task (MWM). In the cue task training, we first tested the ability of the animals to find the platform when it was visible. Under these conditions, the animals from all four groups quickly learned to navigate to the platform and did not show any significant differences in neither the escape latencies nor the average speed, suggesting that vision and navigation properties were not different between the 4 groups (**Suppl. Fig. 1**).

In the subsequent test, the platform was submerged at a different position than previously, and cues were placed on the walls to encourage spatial navigation. The performance across the four training days was analyzed using RM two-way ANOVA. A significant main effect of group (F(3, 272) = 9,202 P<0.0001) and training day (F(3, 816) = 16,16 P<0.0001) was observed without significant main effect of group x training day interaction.

During the four training sessions (day 1-4), the four groups started out on day 1 at slightly different levels, which were however, not significantly different (p>0.49; compare Suppl. Fig. 1D). The WT groups showed a significant learning improvement from training day 1 (D1) to training day 4 (D4; distance travelled to the platform in WT vehicle at D1: 7.36 ± 0.74 m, significantly different from D4: 3.94 ± 0.56 m, Tukey’s p<0.01; WT FTY720: D1: 7.27 ± 0.67 m, and D4: 3.26 ± 0.44 m, Tukey’s p<0.001; **Fig. 3C**).In contrast, the vehicle treated APP/PS1 animals (AD mice) did not improve their learning performance between the first and last day of training (AD Vehicle at D1: 7.18 ± 0.82 m, and D4: 5.86 ± 0.77 m, Tukey’s p=0.52). However, the AD animals receiving chronic fingolimod (FTY720) treatment showed a significant improvement in learning performance, similar to the observation in the WT groups (AD FTY720 at D1: 8.58 ± 0.85 m, and D4: 5.66 ± 0.74 m, Tukey’s p<0.05).

Importantly, when the platform was removed and the animals were tested for long-term memory 24 h later, the RM two-way ANOVA revealed a significant main effect of quadrant (F (1, 65) = 63,63, P<0.0001) with no main effects of group or group x quadrant interaction. Fingolimod (FTY720) treated APP/PS1 animals showed accurate discrimination between the target and the other quadrants, as observed in the WT vehicle and the WT fingolimod groups. In detail, fingolimod treated APP/PS1 mice spent significantly more time in the target quadrant (T) than in the averaged 3 non-target compartments (% time AD FTY720 target: 40.167± 4.105; non-target: 19.94± 1.37,Tukey’s p<0.001, **Fig. 3D**). In contrast, vehicle treated APP/PS1 mice failed to show a clear preference for the target quadrant (% time AD Vehicle target: 38.05 ± 3.43; non-target: 20.65 ± 1.14, Tukey’s p<0.001 **Fig.3D**). As expected, both, vehicle and fingolimod treated WT littermates, occupied the target quadrant significantly longer than the non-target compartments as is also evident from the heat maps shown in Fig. 3E.

Taken together, fingolimod treated APP/PS1 mice showed improved spatial learning in the training session and restored long-term memory in the probe trial compared to vehicle treated APP/PS1 mice, demonstrating that chronic fingolimod treatment rescued the learning impairments in APP/PS1 mice.

### Chronic fingolimod (FTY720) treatment reduces activation of microglia and astrogliosis in APP/PS1 mice

Since fingolimod was shown previously to affect microglia signaling in mouse models of AD (Aytan et al., 2016), we used anti-Iba1 immunohiostological stainings to investigate microglia proliferation/growth, and the extent of microglia activation in brain slices from APP/PS1 and WT littermate mice (**Fig. 4**).

**Fig. 4:**
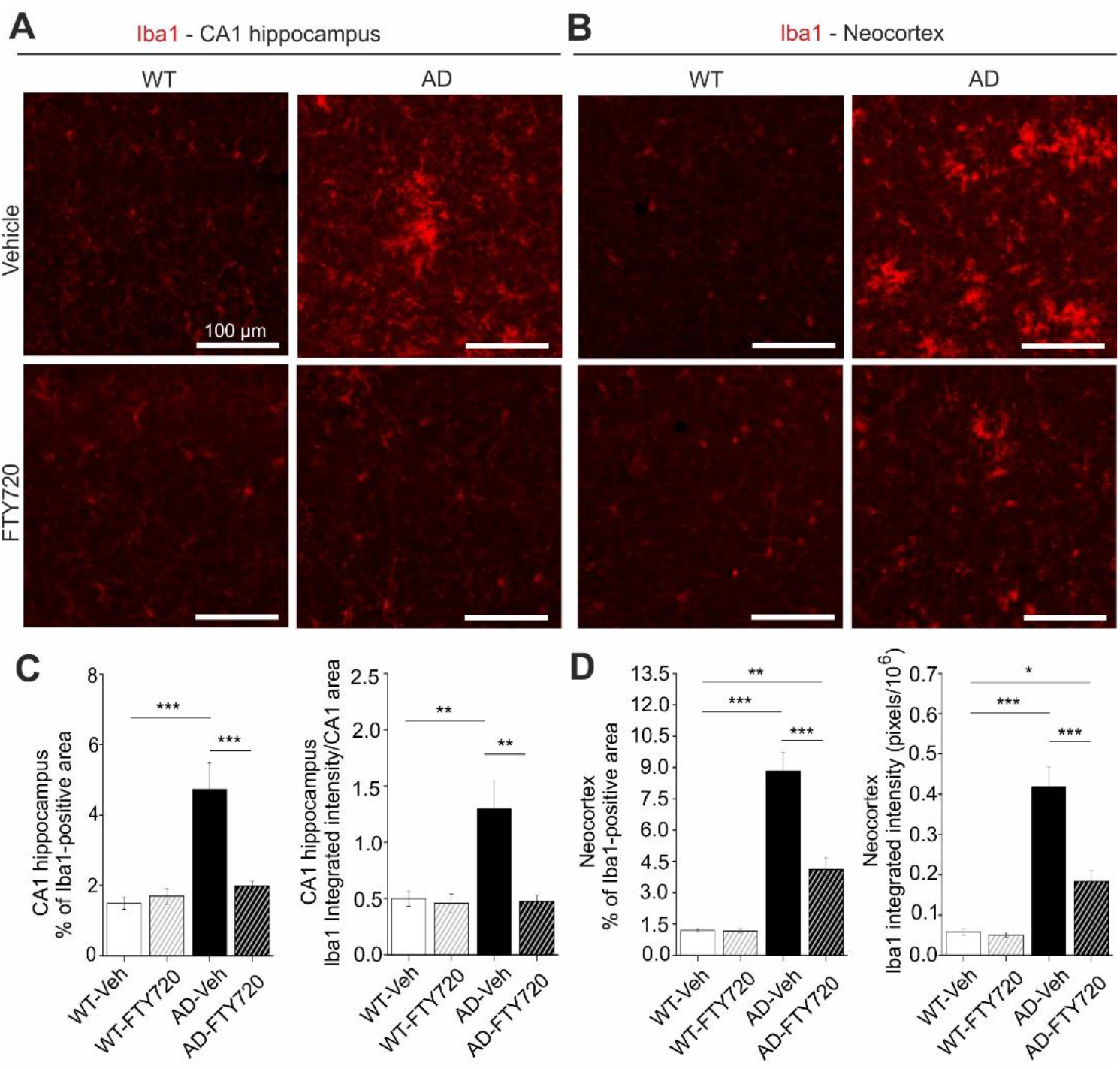
Chronic fingolimod (FTY720) treatment reduces microgliosis in hippocampus and neocortex of APP/PS1 mice. 5-6 months old male WT and APP/PS1 mice were treated with i.p. injections of fingo-limod for 1-2 months. Iba1 immunohistochemistry was performed in the hippocam-pal CA1 area (**A, C**) and in the neocortex (**B, D**) at 6-7 months. **(A)** Iba1 expression was detected with a red fluorescent secondary antibo-dy. Typical examples of Iba1 expression in vehicle and fingolimod treated WT (left) and APP/PS1 mice (right). Illumination and exposure times as well as thresholding of pictures for quantitative analysis were identical for all 4 groups. **(B)** Same analysis as described for (A) but in the neocortex. Typical examples of Iba1 expression in vehicle and fingolimod treated WT (left) and APP/PS1 mice (right). Illumination and exposure times as well as cortical area and thresholding procedures of pictures for quantitative analysis were identical for all 4 groups. **(C)** Quantification of the percent area of CA1 sections as shown in (A) that is covered by Iba1 immune fluorescence (**left**, representing overall microglia area), and of the integrated red fluorescent pixel intensity normalized to the area of the section (**right**, representing the average Iba1 expression). Note the ~3 times increased microglia area and integrated Iba1 intensity in vehicle injected APP/PS1 mice that is completely rescued by fingolimod treatment. Horizontal bars indicate statistical significance between selected groups. **(D)** Same quantification as in (C) but for cortical sections. Vehicle treated APP/PS1 mice displayed increased cortical microgliosis that was significantly reduced upon fingolimod treatment. Significance of differences was tested with two-way ANOVA followed by Tukey’s post hoc test (n=6 animals per WT Vehicle, AD Vehicle and AD +FTY720 groups and 7 animals in WT + FTY720 group). Significance level was set to 0.05 (p<0.05). Different levels of significance are indicated by stars, with * = p<0.05, ** p< 0.01 and *** = p< 0.001.

Here, two-way ANOVA analysis revealed a significant main effect of genotype x treatment interaction (F (1, 21) = 14.46 P=0.0010). In vehicle treated APP/PS1 mice, the percentage of hippocampal CA1 area covered by Iba1 positive microglia was increased threefold compared to WT littermates (AD vehicle: 4.73 ± 0.73%, WT vehicle: 1.48 ± 0.17%), reflecting growth and/or proliferation of microglial cells (**Fig. 4C, left**). Fingolimod (FTY720) treatment (1-2 months, starting at 5-6 months) reduced the area of Iba1-positive microglia back to control levels (AD + FTY720: 1.97 ± 0.14%, WT + FTY720: 1.69 ± 0.22%), indicating complete rescue from microgliosis. The integrated intensity of Iba1 staining divided by the analyzed CA1 area (i.e. normalized Iba1 intensity; **Fig. 4C, right**) is a measure of both, the area covered by microglia and the intensity of Iba1 expression in these cells. Also here, a two-way ANOVA revealed a significant main effect of genotype x treatment interaction (F (1, 21) = 8.565 P=0.0081). The observed 2.5-fold increase in this normalized Iba1 intensity in vehicle treated APP/PS1 mice compared to WT littermates (WT vehicle: 0.50 ± 0.07, WT + FTY720: 0.46 ± 0.08, AD vehicle: 1.30 ± 0.25, AD + FTY720: 0.48 ± 0.05) was similar as compared to the increase in microglia area (3-fold increase; **Fig. 4C, left**). Together these data suggest strongly increased microglial coverage of CA1 in untreated APP/PS1 mice, without an additional effect on the Iba1 expression level per microglial cell area. This finding is corroborated by the similar mean anti-Iba1 fluorescence intensity values observed for all 4 groups (compare **Suppl. Fig. 2**). Fingolimod treatment reduced also the value for normalized Iba1 intensity back to WT control levels, indicating a complete rescue. Together, both types of quantification suggest that fingolimod is a highly effective negative regulator of microglial growth in the CA1 area of the hippocampus. Given the rescue of physiological functions of CA1 pyramidal neurons (spines and LTP) and the rescue of hippocampus-dependent learning (MWM) in response to the fingolimod treatment, our Iba1 results suggest a causal link between the strong anti-inflammatory action of fingolimod on microglia and the rescue of synaptic and memory functions in APP/PS1 mice.

Since cortical dysfunction contributes to reduced memory performance in AD, we also tested microglia activation in the neocortex of our APP/PS1 mouse model. We found that the area of Iba1 covered neocortex and the normalized Iba1 intensity in untreated APP/PS1 mice were both affected more severely than in the hippocampus (**Fig. 4B, D**). Importantly, the fingolimod treatment also dramatically reduced microglia associated neuro-inflammation in this brain area. Two-way ANOVA analysis showed a significant main effect of genotype x treatment interaction for Iba1 coveread area (F (1, 21) = 22.22 P=0.0001), and also for normalized Iba1 intensity (F (1, 21) = 17.42 P=0.0004) in the neocortex.

Interaction of astrocytes and microglia regulate neuroinflammatory processes in neurodegeneration. In the light of the strong functional rescue of memory formation, LTP, and spine density by fingolimod observed in this study, we asked whether astrogliosis was reduced in parallel. Using GFAP IHC we found that the percentage of hippocampal CA1 area covered by astrocytes in vehicle injected APP/PS1 mice was increased roughly twofold (WT vehicle: 4.63 ± 0.33, AD vehicle: 8.01 ± 1.32) compared to vehicle injected WT littermate controls (**Fig. 5**). Strikingly, fingolimod treatment of APP/PS1 mice decreased astrocyte area back to WT control levels (WT + FTY720: 2.73 ± 0.28, AD + FTY720: 2.71 ± 0.37; **Fig. 5C, left**). Likewise, the level of GFAP expression per CA1 area (i.e. normalized GFAP intensity) was regulated in the same way (WT vehicle: 1.51 ± 0.13, WT + FTY720: 0.86 ± 0.16, AD vehicle: 2.86 ± 0.47, AD + FTY720: 0.91 ± 0.15; **Fig. 5C, right**). Two-way ANOVA analysis showed a significant main effect of genotype x treatment interaction for GFAP coveread area (F (1, 21) = 5.999 P=0.0232), and also for normalized GFAP intensity (F (1, 21) = 6.184 P=0.0214). Together, this indicates reduced astrogliosis in the CA1 area of APP/PS1 mice in response to fingolimod treatment. Similar to the results for Iba1, the expression level of GFAP per astrocyte cell area was not different between the 4 groups (**Suppl. Fig. 2**), indicating that GFAP expression levels were not altered by the treatment.

**Fig. 5:**
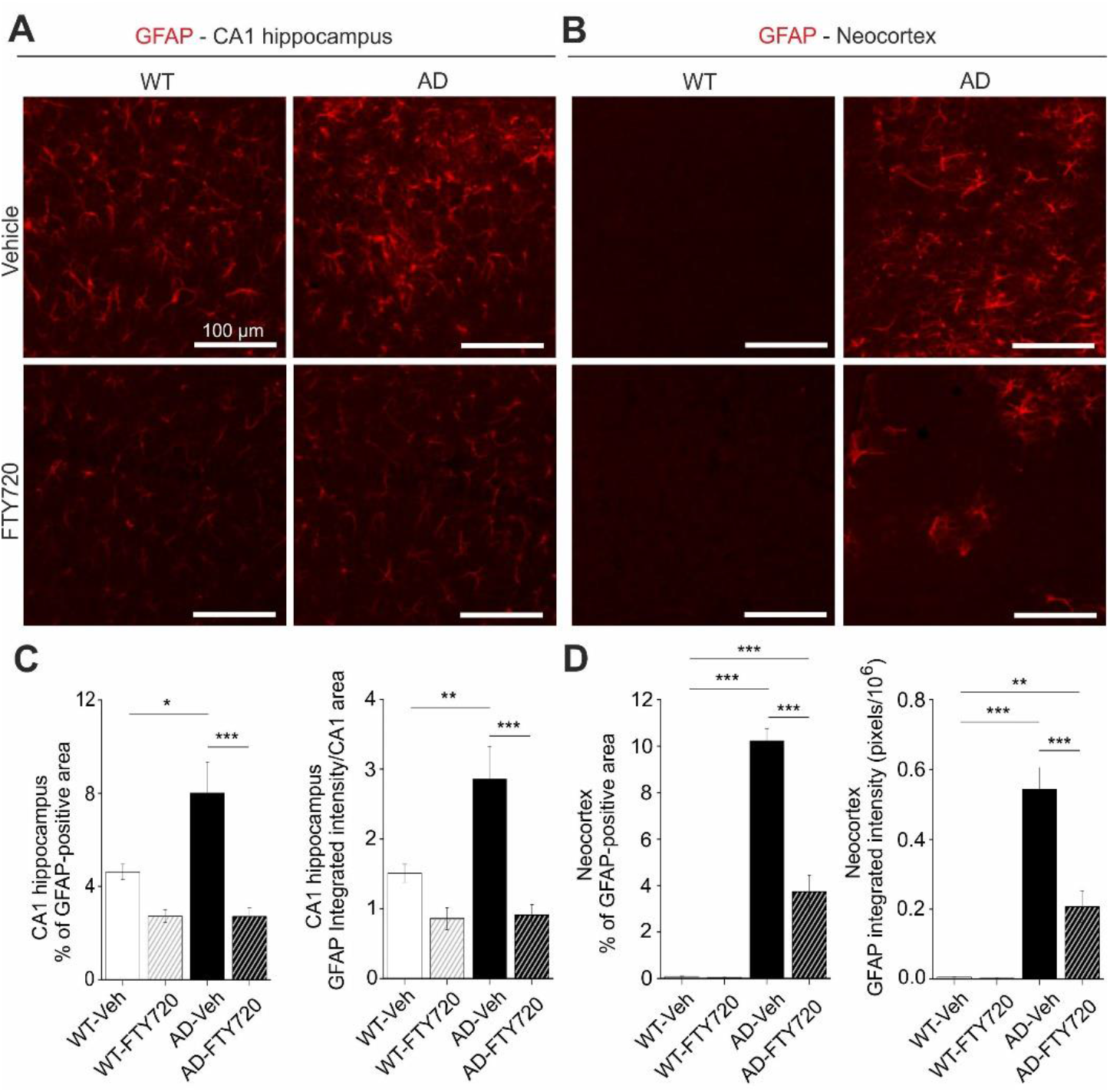
Chronic fingolimod (FTY720) treatment ameliorates astrogliosis in hippocampus and neocortex of APP/PS1 mice. 5-6 months old male WT and APP/PS1 mice were treated with i.p. injections of fingolimod for 1-2 months. GFAP immunohistochemistry was performed in the hippocampal CA1 area (**A, C**) and in the neocortex (**B, D**) at 6-7 months. **(A)** GFAP expression was detected with a red fluorescent secondary antibody. Typical examples of GFAP expression in vehicle and fingolimod treated WT (left) and APP/PS1 mice (right). **(B)** Same analysis as described for (A) but in the cortical sections. Illumination and exposure times as well as thresholding procedures of pictures for quantitative analysis were identical for all 4 groups in (A) and (B). **(C)** Quantification of the percent area of CA1 sections as shown in (A) that is covered by GFAP immunofluorescence (**left**, representing overall area which is covered by activated astrocytes), and of the integrated red fluorescent pixel intensity normalized to the CA1 area in the section (**right**, representing the average GFAP expression). Note the ~2-fold increased area of activated astrocytes and integrated GFAP intensity in vehicle injected APP/PS1 mice that is completely rescued by fingolimod treatment. **(D)** Quantification as described in (C) but for cortical sections. Vehicle treated APP/PS1 mice displayed intense astrogliosis in the neocortex that was significantly reduced upon fingolimod treatment. Horizontal bars indicate statistical significance between selected groups. Significance of differences was tested with two-way ANOVA followed by Tukey’s post hoc test (n=6 animals per WT Vehicle, AD Vehicle and AD +FTY720 groups and 7 animals in WT + FTY720 group). Significance level was set to 0.05 (P<0.05). Different levels of significance are indicated by stars, with * = p<0.05, ** p< 0.01 and *** = p< 0.001.

Of note, fingolimod showed a trend towards reduced astrogliosis also in the hippocampus of WT animals. Although this effect did not reach significance, this suggests that our regime of fingolimod treatment might regulate basal growth of astroglia under control conditions. Similar to Iba1 we also investigated changes in GFAP in the neocortex and found more dramatic astrogliosis in the cortex than in CA1 of untreated APP/PS1 mice, which was effectively reduced by a factor of 2.5 upon fingolimod treatment (**Fig. 5B, D**). Two-way ANOVA analysis showed a significant main effect of genotype x treatment interaction for GFAP covered area (F (1, 21) = 56.75 P<0.0001), and also for normalized GFAP intensity (F (1, 21) = 20.12 P=0.0002) in the neocortex.Overall, astrogliosis and microgliosis in APP/PS1 mice are both significantly reduced by chronic fingolimod treatment and both processes together are likely to account for the rescue of LTP, spine density, and memory formation in our fingolimod treated APP/PS1 mice.

### Chronic fingolimod treatment reduces plaque formation and Aβ deposits in the hippocampus of APP/PS1 mice

Next, we used Thioflavine S staining to investigate changes in the number and size of Aβ plaques in the hippocampus and in the neocortex of APP/PS1 mice (**Fig. 6**). We found that the percent area of the hippocampus that was covered by plaques was reduced roughly twofold by fingolimod treatment, although the effect was not reaching significance (AD vehicle: 0.30 ± 0.11, AD + FTY720: 0.14 ± 0.03; **Fig. 6C**). Smaller and also non-significant decreases upon fingolimod treatment were observed for the number of Aβ plaques per unit area, and for the average size of Aβ plaques (**Fig. 6C**). In the neocortex, plaque load and plaque size were both significantly reduced compared to vehicle treated controls, whereas plaque numbers remained unaffected (**Fig. 6E**). Overall, this analysis revealed a trend towards reduced Aβ plaque burden in hippocampus and a significant reduction in the neocortex.

**Fig. 6:**
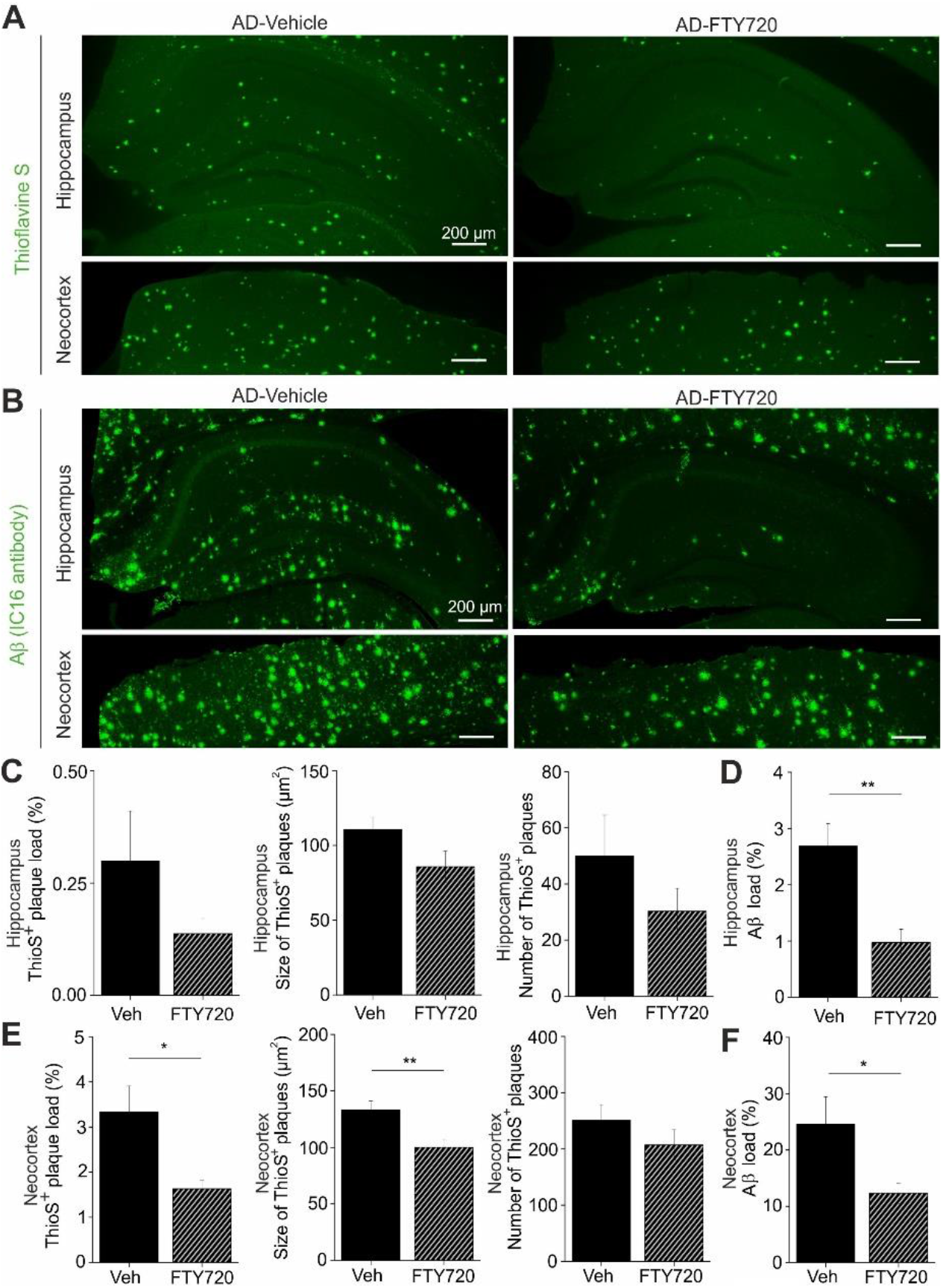
Aβ plaques and other Aβ deposits are reduced in fingolimod (FTY720) treated APP/PS1 mice. 5-6 months old male APP/PS1 mice were treated with i.p. injections of fingolimod for 1-2 months and analyzed for Aβ depositions at 6-7 months. **(A)** Aβ plaques in the hippocampus (upper panels) or in the neocortex (lower panels) were stained with green fluorescent Thioflavine S. **(B)** Overall Aβ protein deposits were detected with an anti-amyloid-β antibody and labeled with a green fluorescent secondary antibody in the hippocampus (upper panels) and in neocortex (lower panels). Representative pictures of Thioflavine S-positive plaques (A) and Aβ depositions (B) are shown for vehicle (left) and fingolimod treated APP/PS1 mice (right). Illumination settings and exposure times as well as thresholding procedures of pictures for quantitative analysis were identical for vehicle and FTY720 treated hippocampal and cortical sections, respectively, for both types of staining. **(C)** Quantification of Thioflavine S-positive amyloid plaque load, expressed as the percentage area of positive staining (left), plaque size (middle), or number of plaques (right) in the hippocampus showed a trend towards reduction in fingolimod treated APP/PS1 mice. **(D)** Interestingly, overall Aβ immunoreactivity in the hippocampus was significantly reduced in fingolimod treated APP/PS1 mice compared to vehicle controls. **(E)** The same quantification as shown in (C) for hippocampus was performed in neocortex and revealed a significantly decreased percentage of plaque load (left) and plaque size (middle) after fingolimod treatment, while the number of plaques (right) displayed only a trend towards reduction in FTY720 treated APP/PS1 mice compared to vehicle controls. **(F)** The percentage of Aβ immunopositive area in the cortex was also significantly reduced in FTY720 treated APP/PS1 mice compared to vehicle controls. Horizontal bars indicate statistical significance between selected groups. Significance of differences was tested with unpaired Student’s t-test (n=6 animals group). Significance level was set to 0.05 (p<0.05). Different levels of significance are indicated by stars, with * = p<0.05, ** p< 0.01.

To get further insight into Aβ protein accumulation in treated and untreated animals, we performed, in slices from the identical individual mice as used in the Thioflavine S experiments, anti-Aβ immunohistological stainings, which detect in addition to Aβ plaques also less condensed Aβ protein deposits (including oligomers and fibrils; **Fig. 6 B**). As expected, this analysis yielded higher values for the percent area covered by Aβ positive material (hippocampus: 2.69 ± 0.40%, neocortex: 24.8 ± 5.0%, **Fig. 6 D, F**) than the Thioflavine S staining in vehicle treated APP/PS1 mice (**Fig. 6C, E**). Following treatment with fingolimod we observed a strong reduction in the percent area covered by Aβ positive structures in both brain areas (**Fig. 6D, F**).

Overall, these data suggest that the fingolimod induced rescue of functional plasticity and neuro-inflammatory signaling in the hippocampus of APP/PS1 mice is accompanied by reduced accumulation of Aβ protein. This reduction of Aβ deposits under fingolimod treatment might be a secondary consequence of the reduced microgliosis we observe, since activated pro-inflammatory microglia was shown recently to contribute to Aβ plaque load via ASC specks formation (Venegas et al., 2017).

Of note, we did not observe any effect of fingolimod treatment on BDNF protein levels or TrkB signaling efficacy in APP/PS1 mice or WT littermates (compare **Suppl. Fig. 3**).

## Discussion

Treatment of Alzheimer’s disease in humans requires medication that can start after onset of disease symptoms. Such therapies are currently not available. Our results show that chronic treatment of APP/PS1 mice with fingolimod (FTY720), starting at 5-6 months - shortly after onset of synaptic and learning dysfunction in these animals - restores functional and structural plasticity as well as learning deficits in this mouse model. This rescuing effect is apparently mediated by reduced microglia mediated neuro-inflammation and reduced astrogliosis, and is accompanied by diminished Aβ accumulation in hippocampus and neocortex. These results demonstrate a promising potential of fingolimod treatment as part of a pharmacological intervention strategy that can ameliorate AD deficits even after onset of the symptomatic phase of the disease.

### Concomitant rescue of deficits in spine density, LTP, and hippocampus-dependent learning in APP/PS1 mice by fingolimod (FTY720)

To the best of our knowledge, this is the first study that reports fingolimod (FTY720) induced combined amelioration of functional and structural synaptic plasticity deficits and of memory dysfunctions in an AD mouse model at 6-7 months of age with fingolimod treatment starting no earlier than at 5 months. Interestingly, a recent study by Carreras et al. (Carreras et al., 2019) reported prevention of Aβ pathology and of memory deficits in the 5xFAD Alzheimer mouse model when treatment commenced at 1 month of age and was continued until start of experiments at 8 months. Our results go beyond this previous study by showing reversal of spine, LTP, and memory deficits in response to 1-2 months of fingolimod treatment in an APP/PS1 AD mouse model when started after outbreak of disease symptoms.

The spine deficits observed at this age in untreated AD mice strongly depended on the proximity of CA1 pyramidal cell dendrites to the closest Aβ plaque. This observation was consistent with the previously reported spine instability recorded *in vivo* in the neocortex of 3-4 months old animals of the same APP/PS1 mouse model as used in our study, which was also more pronounced in the vicinity of Aβ plaques (Liebscher et al., 2014). These results suggest that increased instability of spines in living tissue (Liebscher et al., 2014) correlates with net spine loss that can be detected with Golgi-Cox staining (our study). The finding that the spine loss depended on the proximity of spines to plaques is consistent with the idea that soluble monomeric or oligomeric Aβ species, which are at chemical equilibrium with Aβ plaques and accordingly will be present at higher concentrations nearby plaques, are likely the toxic agents that induce spine loss. This seems to corroborate the general assumption that soluble species of Aβ are responsible for synaptic deficits in AD (Mucke and Selkoe, 2012). However, the strong correlation between the degree of microgliosis and synaptic as well as memory dysfunctions in our APP/PS1 mice leads us to suggest that the dependence of spine deficits on proximity to plaques might be related rather to neuroinflammatory signaling of microglia – a cell type that is always closely associated with Aβ plaques. The pronounced rescue of spine deficits in APP/PS1 mice by fingolimod apparently depended on the distance to Aβ plaques and associated microglia (**Fig. 2B**). While spine densities near plaques were also dramatically recovered by fingolimod, they did not reach WT levels, as achieved for spine density more distant to plaques. Since Aβ protein deposits – although significantly decreased – did not vanish under fingolimod treatment, the remaining Aβ oligomers in treated APP/PS1 mice still have disadvantageous effects for spine stability and therefore a complete rescue could not be expected. In this respect, the incomplete spine rescue near plaques strengthens the claim of a causal connection between Aβ pathology and intact spine function. However, the percent increase in spine density relative to vehicle treated AD mice was comparable near and distant from plaques. Thus, future studies using e.g. longer FTY720 interventions are required to resolve whether spine densities near to plaques can be returned to WT levels when using different treatment regimes.

LTP recordings performed in identically treated 6-7 months old APP/PS1 mice and WT littermates revealed strongly impaired LTP magnitudes in untreated APP/PS1 mice compared to WT littermates (**Fig. 3**), whereas basal excitability and synaptic transmission properties were not significantly affected (data not shown). The fingolimod treatment of APP/PS1 mice led to an equally compelling rescue of LTP back to WT control levels as observed for spines distant to Aβ plaques. Since field potential LTP recordings, as used here, read out synaptic efficacy of glutamatergic synapses across the apical dendritic tree of a cell, it is conceivable that after fingolimod treatment most recorded synapses are located distant to plaques. The observed complete rescue of LTP therefore fits well to complete rescue of spines distant to plaques. It remains to be shown whether activity-dependent spine formation is rescued because of reinstated LTP, or rather LTP is rescued because of the higher number of functional spines. Unexpectedly, fingolimod treatment showed a tendency to significantly reduce LTP magnitude in WT animals, whereas no such phenomenon was observed for spines in WT animals. Since WT and APP/PS1 mice differ by the presence of Aβ oligomers, plaques, and associated activated cells (astrocytes and microglia), it could be argued that in APP/PS1 mice fingolimod was scavenged by plaques and associated cells, whereas in WT animals the full concentration of fingolimod was present close to neuronal cell membranes. This higher local concentration could thereby cause impaired synaptic properties in a way that LTP was reduced. Nevertheless, we did not perform additional experiments to address these questions since treatment of healthy animals (or humans) with fingolimod would never be intended.

Spine and LTP integrity are necessary for normal function of learning and memory processes. Concomitant with spines and LTP rescue by fingolimod, we show here that early memory deficits in this AD model are also completely rescued by this treatment. To our knowledge, this is the first evidence suggesting that fingolimod has the potential to rescue cognitive deficits of APP/PS1 mice even when treatment starts after disease onset. At present, as for spines and LTP data, we can only correlate this rescue to lower A⍰ load and lower neuro-inflammation. Either of these events could be responsible for the rescuing action of fingolimod on learning performance since both, increased brain A⍰ and increased levels microglia-derived cytokines, were previously shown to perturb memory processing (Donzis and Tronson, 2014; Mucke and Selkoe, 2012).

Whereas we observed the compelling rescue of AD associated deficits in synaptic plasticity and memory dysfunction by fingolimod treatment (**Figs. 2, 3**), this rescue was not accompanied by BDNF elevations or enhanced TrkB signaling, neither in hippocampus nor neocortex (compare **Suppl**. **Fig. 3**), albeit fingolimod mediated amelioration in other mouse models of brain diseases was associated with increased BDNF/TrkB signaling (see e.g., (Deogracias et al., 2012; Miguez et al., 2015)). However, in our AD mouse model, Aβ pathology develops as a chronic, progressive accumulation of Aβ deposits and our mice were tested for BDNF elevation at 6-7 months after treatment with fingolimod for 1-2 months. Therefore, it seems possible that fingolimod induced BDNF elevations might critically depend on the starting age as well as the schedule and duration of fingolimod application. Interestingly, BDNF levels in our 6-7 months old untreated APP/PS1 mice were not reduced compared to WT littermates (**Suppl. Fig. 3**), suggesting that the cognitive deficits we observed were not causally connected to BDNF reduction in untreated APP/PS1 mice.

### The role of Aβ reduction in fingolimod-dependent rescue of APP/PS1 mouse deficits

Our results obtained with Thioflavine S staining and anti-Aβ immunohistochemistry (**Fig. 6**) suggest that the fingolimod mediated rescue of hippocampal function in APP/PS1 mice was paralleled by reduced Aβ accumulation. Especially when focusing on additional non-plaque Aβ deposits that can be detected with Aβ immunohistochemistry, rather than on Thioflavine S-positive plaques, this reduction in Aβ accumulation was obvious. This result is consistent with previous findings in another AD mouse model (5xFAD), where chronic fingolimod treatment started already in 4 weeks old, non-symptomatic, female animals (Aytan et al., 2016). Nonetheless, neither these related previous results in juvenile mice nor our own data in symptomatic adult AD mice allow to conclude that there is a causal connection between reduced Aβ deposits and rescued hippocampal function in our APP/PS1 mice. However, Thioflavine S plaques are surrounded by Iba1 positive microglia (data not shown). As microglial growth in CA1 is dramatically reduced in response to our fingolimod treatment regime, these data suggest an instrumental role of reduced microglia mediated neuro-inflammation in the rescue of hippocampal functions in fingolimod treated APP/PS1 mice. Since rescue of spine deficits appeared to be more complete distant than close to plaques (compare **Fig. 2B**) this rescuing effect might be associated with reduced Aβ levels or altered microglia signaling (e.g. released IL-1, IL-6, TNF-α). In addition, reduced levels of microglia-derived ASC specks proteins, which have recently been shown to increase Aβ plaque formation (Venegas et al., 2017), might contribute to fingolimod induced rescue of synaptic functions in APP/PS1 mice.

### The role of neuro-inflammation in fingolimod-dependent rescue of APP/PS1 mouse deficits

We observed spine, LTP, and memory rescue in APP/PS1 mice when fingolimod treatment started at the onset of the symptomatic phase of AD pathology. This rescue did not depend on enhanced BDNF protein expression or enhanced downstream TrkB signaling. Since our IHC data showed a strong reduction of astrogliosis and microgliosis in response to treatment with fingolimod, we suggest that amelioration of the destructive neuroinflammatory signaling by both cell types forms the basis to explain the strong rescuing effect of fingolimod.

Fingolimod has complex and only partially resolved signaling functions in the brain, with prominent effects particularly on microglia (reviewed in (Angelopoulou and Piperi, 2019; Hunter et al., 2016)). Fingolimod can pass the blood-brain barrier (BBB) and is partially converted to fingolimod-phosphate (fingolimod-P) in brain cells through the action of sphingosine kinases (SPHK_1, 2_). Both fingolimod species can be released into the extracellular space and bind either as agonist or as modulator to 7-TM spanning G-protein coupled sphingosine-1-phosphate receptors (S1PR_1-5_; compare (Angelopoulou and Piperi, 2019)). Fingolimod induced activation of microglial S1PRs was shown to interfere with microglia activation in the cortex of 5xFAD mice (Aytan et al., 2016), to inhibit neuronal Aβ production (Takasugi et al., 2013), to decrease Aβ_42_ and Aβ_40_ levels in the brain of AD mouse models (Aytan et al., 2016; McManus et al., 2017), to regulate Aβ traffic across the BBB (Sanabria-Castro et al., 2017), and to promote the conversion of pro-inflammatory M1 microglia to the anti-inflammatory M2 state (Qin et al., 2017). With respect to astroglia, fingolimod treatment was shown recently to reduce astrocyte activation and to increase Aβ phagocytosis by astrocytes (McManus et al., 2017). All these beneficial effects of fingolimod against microgliosis and astrogliosis are consistent with the recovery of hippocampal functions of APP/PS1 mice described here. However, these studies can also not resolve whether the dramatic rescue effect of fingolimod in our mouse model is a direct effect of reduced Aβ levels, altered cytokine release from microglia and astrocytes, or related effects leading to microglia-dependent synaptic pruning (Wilton et al., 2019).

Interestingly, also in WT mice FTY720 treatment showed a trend to reduce astrogliosis in CA1 (**Fig. 5C**) and to reduce LTP (**Fig. 3A**). Since astrocyte activity is an important co-factor in hippocampal synaptic plasticity (see e.g., (Nishiyama et al., 2002; Perea and Araque, 2007)), future studies should address whether these FTY720 effects in WT mice are related.

Overall, our results suggest that repurposing of FDA approved anti-inflammatory drugs, such as fingolimod, might be a promising strategy to generate valuable pharmacological tools to treat AD even after disease onset. We provide evidence that such anti-neuroinflammatory drugs can counteract microglia and astrocyte mediated processes that lead to synaptic dysfunction, spine deficits and eventually memory decline. This anti-neuroinflammatory approach should be combined with treatments that elevate neuroprotective growth factors like BDNF (Choi et al., 2018) and other drugs (to be developed) that specifically tackle tau pathology associated dysfunction in the neocortex. Together these drugs might lay the basis for a multi-target combination therapy against AD.

## Materials and methods

### Animals

For all experiments 5-7 months old heterozygous APP/PS1 mice and their wildtype littermates were used. These APP/PS1 mice harbor a mutation in the APP gene (KM670/671NL, “Swedish mutation”) as well as a mutated presenilin 1 (Leu166Pro mutation), both under control of the Thy1 promotor that drives neuron specific expression of both proteins (Radde et al., 2006). Both gene mutations are associated with early-onset familial AD (FAD) in humans. The animals were generated on a C57BL/6J genetic background and constantly backcrossed with C57BL/6J mice (Charles River, Sulzfeld, Germany). The animals were housed in groups of 3 -4 animals and had constant access to food and water. They were maintained on a 12:12 h light dark cycle (lights on at 7 a.m.). All experiments were performed during the light period of the animals and were in accordance with the European Committees Council Directive (2010/63/EU) and were approved by the local animal care committees.

### Fingolimod (FTY720) administration

Male WT and AD transgenic mice were treated with fingolimod (FTY720) at the age of 5-6 months (age range of animals at the start of treatment: 22-24 weeks). The drug was dissolved in 3% DMSO in saline, and was administered every second day via i.p. injection at 1 mg/kg body weight. All animals were treated for 1-2 months according to this scheme until they were sacrificed for electrophysiological, spine, or immunohistological experiments, or until completion of behavioral testing. Electrophysiological experiments, spine analysis, and behavioral testing were done separately in distinct cohorts of animals. Control mice were treated identically with vehicle solution.

### Combined Aβ plaque staining and Golgi-Cox impregnation

All mice were injected twice (24h interval) i.p. with 75 μl of 10 mg/ml Methoxy-X04 (TOCRIS) in DMSO (Sigma-Aldrich) as described previously (Jahrling et al., 2015; Liebscher et al., 2014). Two hours after the second injection, they were anesthetized and transcardially perfused with 0.9% saline followed by 4% PFA/PB (pH 7.4). The injection steps were performed according to Jahrling et al. (Jahrling et al., 2015). The brains were postfixed in 4% PFA/PB pH,7.4 for 24 hours at 4 °C and transferred into Golgi-Cox solution in the dark and the solution was changed only once after 7 days. After 14 days, the brains were placed into 25% sucrose in PBS at 4 °C for at least 1-2 days. Coronal sections of 100 μm thickness were cut using a Vibratome (Pelco Model 1000, The Vibratome Company, St. Louis, USA). Sections were mounted onto gelatin-coated slides and after allowing them to dry, they were washed with distilled water for 2 min and then they were transferred into 20% ammonium hydroxide in distilled water (Ammonium hydroxide solution, ACS reagent, 28.0 – 30.0 % NH3 basis, Sigma-Aldrich). The sections were washed again with distilled water twice for 2 min each. The sections were dehydrated passing through ascending grades of ethanol 70%, 95% and 100% for 5 min each and cleaned in Xylol (Roth) twice for 10 min each. The sections were coverslipped with DePex medium (Serva) and after letting them dry for 2 days they stored at 4 °C ready for analysis. The composition for the Golgi-Cox solution was 5% potassium dichromate (Merck), 5% mercuric chloride (Merck) and 5% potassium chromate (Merck; compare (Das et al., 2013)). All stock solutions as well as the final Golgi-Cox solution were prepared according to the protocol used by Bayram-Weston (Bayram-Weston et al., 2016).

### Image processing and dendritic spine density analysis

Spine density of secondary apical dendritic segments in stratum radiatum (SR) of CA1 pyramidal neurons was calculated as the number of spines per micrometer dendritic length. Ten dendritic segments (15 μm long) per animal were analyzed for all treated and untreated groups. Dendritic lengths were estimated using the NeuronJ plugin of NIH ImageJ software (https://imagej.nih.gov/ij/). For AD mice, 10 dendritic segments which were at a distance more than 50 μm from the plaque border (AD distant) and 10 dendritic segments which were located within 50 μm from the plaque border (AD near) were analyzed per animal. This digital near/distant classification scheme was selected to facilitate quantification of the results. When plotting spine densities versus plaque distance for individual dendritic segments we did not observe a clear cut-off at a certain distance, but a decrease at distances <50 μm is evident (compare **Suppl. Fig. 4**). Images for Golgi-Cox stained dendritic segments and the blue fluorescent methoxy-X04 stained Aβ plaques were captured with a 40x magnification objective using a SPOT digital camera that was attached to a LEITZ DM R microscope (Leica). The distance in z-direction of analyzed dendrites to the border of the Methoxy-X04 stained Aβ plaque was in all cases <5 μm. This deviation was minor compared to the calculated distances in the x-y space and was therefore neglected. Imaging for Golgi-Cox staining was performed in bright field mode while Methoxy-X04 fluorescence was captured with a filter cube (excitation filter: BP 360/40 nm, dichroic mirror: 400 nm, emission filter: LP 425; Leica). The distance of the analyzed dendritic segments from the Aβ plaques was determined as the average distance of both segment end-points and the center of the segment to the respective plaque border in merged pictures using ImageJ software. Dendritic segments from different neurons were traced and spine density was quantified as the number of spines per dendritic length in μm. The spines were counted manually using a 100x magnification oil immersion objective. Spine density analysis was conducted blindly to the treatment in WT and AD mice.

### Preparation of slices for IHC

Coronal slices were prepared from young adult male APP/PS1 mice or WT littermates at an age of 6-7 months. Animals were anesthetized using intraperitoneal injection of Ketamine hydrochloride/Xylazine hydrochloride (Sigma-Aldrich) and then transcardially perfused with ice-cold 0.9% NaCl solution. After opening the cranium, the brains were gently removed and the right hemi-brains were post-fixed in 4% PFA/0.1 M phosphate buffered saline (PBS) for 24 hours at 4 °C and then cryoprotected in 25% sucrose (AppliChem) in PBS. The left hemi-brains were quickly placed in ice-cold ACSF solution, dissected at the anterior and posterior cortex and hippocampus and all brain areas were snap-frozen for protein analysis. The fixed hemispheres were then cut to 40 μm thick coronal slices using a cryostat (Leica CM 3050). Three free-floating sections 240 μm apart from each other (between Bregma levels −2.46 mm and −3.08 mm according to Franklin and Paxinos (Franklin K., 1996), containing the hippocampus and cortex were selected for all histological fluorescent stainings. For imaging experiments, all sections were transferred to slides (Superfrost, Thermo Scientific), coverslipped with ImmunoMount mounting medium (Thermo Scientific).

### *Immunofluorescence for total A*β

Free-floating sections were permeabilized with PBS/0.1 % Triton X-100 and incubated with 98 % formic acid to perform antigen retrieval. This step was followed by washes with PBS. Sections were then blocked in 20 % normal goat serum (Dianova) in PBS/Triton X-100 0.1 % and incubated overnight at 4°C with the mouse anti-Aβ IC16 antibody (1:400; kindly provided by Prof. Claus Pietrzik, Johannes-Gutenberg-University Mainz (Richter et al., 2010) in 10 % normal goat serum (Dianova) in PBS containing 0.1 %Triton X-100. Sections were then washed with PBS/0.1 %Triton X-100, incubated with an Alexa Fluor 488 labeled goat anti-mouse antibody (1:500, Thermo Scientific) in 10 % normal goat serum (Dianova) in PBS/0.1 % Triton X-100, and finally washed with PBS. Imaging was performed with ZEN 2010 software using a 5x objective in a confocal imaging system (LSM 780, Zeiss, Germany). Green fluorescence was excited using the 488 nm laser line from an Argon laser. For Aβ load quantification 16 bit images were converted to 8 bit gray-scale images and after defining the region of interest (total hippocampal or cortical area) they were thresholded within a linear range using the NIH ImageJ software. The load was expressed as the percentage area covered by Aβ-positive staining (% Aβ load). The quantification performed blind to the treatment for WT and AD mice. The quantification was performed blind to the treatment for WT and AD mice.

### Immunofluorescence for GFAP

For GFAP staining, antigen retrieval was performed with sodium citrate buffer (10 mM, pH 6.0) for 30 min at 80 °C followed by 3 washes with TBS. Sections were blocked in 5% normal goat serum (Dianova) in TBS with 0.4 % Triton X-100 and incubated with a mouse anti-GFAP antibody (1:500, clone G-A-G, Sigma: G 3893) in blocking solution overnight at 4°C. Sections were washed with TBS and incubated with an Alexa Fluor 488 labeled goat anti-mouse antibody (1:500,Thermo Scientific) in blocking solution and were finally washed with TBS. For the quantification of GFAP in the CA1 area of the hippocampus and cortex images were captured with a 10x objective using a SPOT camera attached to a Leica LEITZ DM R microscope through an appropriate filter cube (excitation filter: BP 515-560 nm, dichroic mirror: 580 nm, emission filter: LP 590 nm, Leica). Using NIH ImageJ software for quantification, the load was expressed as the percentage area covered by GFAP-positive staining and the normalized integrated intensity (raw integrated density divided by the CA1 area) was expressed in pixels (pixels/106). The quantification was performed blind to the treatment for WT and AD mice.

### Immunofluorescence for Iba1

Free-floating sections were first treated with sodium citrate buffer (10 mM, pH 6.0) for 30 min at 80°C for antigen retrieval and then washed 3 times with PBS. Sections were blocked in 10 % FBS (Gibco) and 1 % BSA (Sigma) in PBS containing 0.3 % Triton X-100 and incubated overnight at 4°C with a rabbit anti-Iba1 antibody (1:500, Wako) in PBS containing 1 % FBS (Gibco), 0.1 % BSA (Sigma) and 0.3 % Triton X-100. Sections were then washed with PBS and developed with an Alexa Fluor 555 labeled donkey anti-rabbit antibody (1:500, Thermo Scientific) in PBS containing 1 % FBS, 1 % BSA, and 0.3 % Triton X-100. The sections were finally washed with PBS. Imaging and quantification were performed as described for GFAP immunohistochemistry. The quantification was performed blind to the treatment for WT and AD mice.

### Thioflavine S staining

Coronal free-floating sections were incubated for 9 min in 1% Thioflavine S (Sigma-Aldrich) aqueous solution and then treated with 80% ethanol 2 times for 3 min each, followed by a wash with 95% ethanol for 3 min. Sections were rinsed 3 times with distilled water. Thioflavine S imaging was performed with a confocal Zeiss laser-scanning-microscope using Zen software (LSM 780, Zeiss, Germany). A 5x magnification objective was used and green fluorescence was excited with a 488 nm Argon laser. Primary images were converted to 8 bit gray-scale and after defining the region of interest (total hippocampal or cortical area) they were thresholded using NIH ImageJ software. Thioflavine S plaque load was expressed as the percentage area covered by Thioflavine S-positive staining (% ThioS). The ImageJ tool ‘Analyze particles’ was used to determine plaque number and size (plaque area in μm^2^). The quantification was performed blind to the treatment for WT and AD mice.

### Western blotting

The hippocampal brain samples from WT and APP/PS1 mice chronically treated with fingolimod or vehicle were homogenized using ultrasound sonicator in NP lysis buffer containing 2% orthovanadate and 4% protease inhibitor mix. The homogenized suspension was centrifuged (15,000 g, 10 min, +4°C), and the resulting supernatant was used for analysis. Proteins were separated with SDS-PAGE and transferred to PVDF membrane. The membranes were incubated with antibodies against phosphorylated Trk/TrkB at residues Y515, (C35G9, Cell Signaling Technology, #4619), Y706 (Cell Signaling Technology, #4621), Y816 (Cell Signaling Technology, #4168), total TrkB (R&D Systems, # AF1494), phospho-p70 S6 Kinase (Cell Signaling Technology, #9205), total p70 S6 Kinase (Cell Signaling Technology, #9202), and β-Actin (Sigma Aldrich, #A1978). Following incubation with primary antibodies, the membranes were incubated with horseradish peroxidase conjugated secondary antibodies, and the bands were visualized using chemiluminescent western blotting substrate with a Fuji LAS3000 camera (Tamro Medlabs, Finland). Phosphoproteins were normalized against total proteins, total proteins were normalized against β-Actin.

### BDNF quantification

To assess the levels of hippocampal BDNF protein the BDNF Quantikine ELISA kit (R&D Systems, Wiesbaden, Germany) was used. Total hippocampi were dissected and processed according to the manufacturer’s instructions (compare (Petzold et al., 2015)). Protein levels were set in relation to the wet weight of the brain tissue sample.

### Preparation and LTP recordings of acute hippocampal slices

Acute transverse hippocampal slices (350 μm thick for field recordings) were obtained from APP/PS1 and WT mice. Animals were first anaesthetized with isoflurane and killed in accordance with the European Communities Council Directive (80/609/EEC). Slices were cut on a vibratome (Microm HM600V, Thermo Scientific) in ice-cold dissecting solution containing (in mM): 234 sucrose, 2.5 KCl, 0.5 CaCl2, 10 MgCl2, 26 NaHCO3, 1.25 NaH2PO4 and 11 D-glucose, oxygenated with 95% O2 and 5% CO2, pH 7.4. Slices were first incubated, for 60 min at 37°C, in an artificial CSF (ACSF) solution containing (in mM): 119 NaCl, 2.5 KCl, 1.25 NaH2PO4, 26 NaHCO3, 1.3 MgSO4, 2.5 CaCl2 and 11 D-glucose, oxygenated with 95% O2 and 5% CO2, pH 7.4. Slices were used after recovering for another 30 min at room temperature. For all experiments, slices were perfused with the oxygenated ACSF at 31 ± 1 °C. Field EPSPs were recorded in the stratum radiatum of the CA1 region using a glass electrode (RE) (filled with 1 M NaCl, 10 mM HEPES, pH 7.4) and the stimuli (30% of maximal fEPSP) were delivered to the Schaeffer Collateral pathway by a monopolar glass electrode (SE) (filled with ACSF). The recordings were performed using a Multiclamp 700B (Molecular Devices) amplifier, under the control of pClamp10 software (Molecular Devices).

LTP was induced using a high-frequency stimulation protocol (HFS) protocol with two pulses of 100 Hz spaced by 20 seconds inter-stimulus interval. For LTP analysis, the first third of the fEPSP slope was calculated in baseline condition (20 minutes prior to induction protocol delivery and for 45-60 minutes post-induction). The average baseline value was normalized to 100% and all values of the experiment were normalized to this baseline average (one minute bins). The recordings were performed blind to the treatment for WT and AD mice.

### Morris water maze

Spatial learning and memory were tested with a Morris water maze paradigm. The apparatus consisted of a circular tank (Ø 90 cm) filled with water (temperature 23 ± 1 °C) made opaque with the addition of 100 ml of white opacifier (Viewpoint, France). The test consisted of 3 phases: (1) Cue task training (2 days), (2) Spatial learning training (4 days) and (3) Long-term reference memory-probe test (1 day). Cued task was done prior to spatial learning training to detect visual and motor problems and to accustom the mice to the testing rule (find the platform to escape). A visible flag was placed on top of the platform and the maze was surrounded by opaque curtains. The mice were allowed to find the visible platform. The platform position was changed for the second cue task (day2). The escape latency, average speed and distance travelled were recorded. For the spatial learning, the extra-maze cues were mounted on the side-walls. A new platform position was chosen and kept in the same position for the 4 training days. If the animal did not find the platform within the trial duration it was gently guided to it. For the cue task and spatial trainings, all animals performed 4 trials/day with a maximum trial duration of 90 s (+30 s on the platform at the end of each trial) and an inter-trial interval of 10 min. Twenty-four hours after training completion, the platform was removed and a probe test was run for 60 s. All the trials were video recorded and tracked using ANYmaze software. The distance travelled to reach the platform for each trial was analyzed. During the probe test, the distance travelled in each quadrant (target (previous position of the platform), left, right and opposite) was recorded and analyzed.

### Statistical analysis

Statistical analysis was performed by using GraphPad Prism (GraphPad Software Inc., La Jolla CA, USA) software. Spine densities and quantification of IHC stainings were analyzed either by two-sided t-test comparisons or, in case of multiple comparisons, by one-way or two-way ANOVA (analysis of variance), as indicated. For LTP and behavioral analyses, two-way ANOVA or repeated measures (RM) two-way ANOVA was applied followed by Tukey’s post-hoc analysis when appropriate. All data were checked for normal distribution by using the Shapiro-Wilk test. If not otherwise indicated, all data followed the normal distribution. Statistical significance was determined as p<0.05. All data are depicted as mean ± standard error of the mean (SEM).

## Declarations

### Ethics approval

All experiments were performed in accordance with the European Committees Council Directive (2010/63/EU) and were approved by the local animal care committees.

### Consent for publication

All authors have approved the final version of the manuscript and support submission in its current form.

### Availability of data and material

All data and material will be made available to other scientist upon appropriate requests.

### Authors’ contributions

Experiments were performed by: G-IK (spine analyses, all immunohistochemical procedures, all fluorescence microscopy data; Figs. 2, 4, 5, 6 and Suppl. Fig. 2); ARSP (behavioral analysis; Fig. 3, Suppl. Fig. 1); TE (planning of animal treatments, BDNF ELISA analysis; Fig. 1 and Suppl. Fig. 3); PP and KD (LTP recordings; Fig. 3, Suppl. Fig. 1); AL and PC (TrkB signaling; Suppl. Fig. 3); EE (characterization of APP/PS1 mice). Experiments were designed by G-IK, TE, PP, EE, HM, EC, KG, and VL. The data were analyzed by G-IK, ARSP, PP, TE, and AL. The study was designed and supervised by KG, HM, TE, and VL. The manuscript was written by VL, G-IK, and TE with help from KG, EC, and HM. VL and KG initiated and conceived the project.

## Acknowledgements

The authors want to thank Dr. Thomas Munsch for help with fluorescence and confocal microscopy, Sascha Weggen for advising Aβ quantification, Jan Dreßler and Anja Reupsch for processing of histological sections, and Margit Schmidt for ELISA analysis.

