## Supplemental material for "Anti-inflammatory treatment with FTY720 starting after onset of symptoms reverses synaptic and memory deficits in an AD mouse model"

### Supplemental Materials

#### Supplemental Results

**Fingolimod (FTY720) treatment does not affect BDNF expression or TrkB signaling in APP/PS1 mice**

Fingolimod treatment of mice was reported previously to increase BDNF protein expression in the hippocampus of mouse models of different neurodegenerative diseases (Deogracias et al., 2012; Fukumoto et al., 2014), whereas others observed ameliorating effects of fingolimod in a mouse model of Huntington's disease in the absence of BDNF protein elevation (Miguez et al., 2015). When testing BDNF protein levels in hippocampus and neocortex of our fingolimod treated mice with ELISA (**Suppl. Fig. 3A**), we did not observe any changes in response to fingolimod treatment, neither in APP/PS1 mice nor in WT littermates. Interestingly, we found roughly 30% increased BDNF protein levels in the neocortex of APP/PS1 mice compared to WT control, being consistent with a similar compensatory increase in BDNF levels previously described in another APP/PS1 mouse model (Rantamaki et al., 2013). However, also this increase of BDNF in the neocortex was not further enhanced by fingolimod treatment. While these data indicate that our ELISA approach was sensitive enough to detect the compensatory BDNF elevation in the cortex of APP/PS1 mice, the absence of any effect of fingolimod on BDNF protein expression in cortex and hippocampus suggests that the fingolimod induced rescuing effects of neuronal function we observe here are not mediated by an elevation of BDNF protein expression in hippocampal neurons.

To determine whether fingolimod affects signaling of released BDNF via TrkB receptors we analyzed tyrosine phosphorylation of TrkB and activation of TrkB downstream signaling cascades. Here, we observed a significantly reduced phosphorylation of Y816 of TrkB and a trend towards reduced phosphorylation of Y515 and Y706 upon chronic fingolimod treatment in both, APP/PS1 and WT animals, while overall TrkB levels remained unaffected by (**Suppl. Fig. 3B**). Overall TrkB protein levels were not significantly different between WT and APP/PS1 mice (data not shown). From the set of TrkB downstream signaling partners tested, only phosphorylation of the p70 S6 kinase was slightly decreased upon fingolimod, albeit not reaching significance (data not shown). Thus, we did not observe a similar increase in TrkB signaling cascades in response to fingolimod treatment, as it was previously described for other mouse models of neurodegenerative diseases (Deogracias et al., 2012; Miguez et al., 2015). If anything, fingolimod decreased TrkB signaling. Taken together, these biochemical results speak against any elevation of TrkB-dependent signaling during chronic fingolimod treatment, corroborating the lack of an effect of fingolimod also on hippocampal BDNF protein levels.

#### Supplemental Figures

**Suppl. Fig. 1: Absence of any change in synaptic fatigue during high frequency stimulation LTP induction, in post-tetanic potentiation, and in Cue task motor parameters during Morris Water Maze experiments upon fingolimod (FTY720) treatment.**

Same experimental conditions as in Fig.3. **(A)** Peak fEPSP amplitudes of the first 15 pulses of the first tetanic induction train were analyzed in the four groups; **(B)** The first minute of the fEPSP slope post-induction, representing post-tetanic potentiation, was measured in all four groups (WT-vehicle n=11; WT-FTY720 n=8; AD Vehicle n=8; AD FTY720 n=8). **(C)** Average speed (m/s) and distance travelled measured during days 1 and 2 (D1, D2) of the cue task of WT Veh (n=18), AD Veh (n=15), WT FTY720 (n=20) and AD FTY720 (n=16) mice. n=number of animals. **(D)** Average performance in the MWM of the 4 groups of animals in trials (t) 1-4 on training day 1. Two-way ANOVA revealed absence of a significant difference between the 4 groups.

**Suppl. Fig. 2: Unchanged mean fluorescence intensity per area of microglial and astroglial cells in hippocampus and cortex upon fingolimod (FTY720) treatment**

Same experimental conditions as in Figs.4 and 5. The graphs show the mean fluorescence intensity value per pixel of microglia and astrocyte area in hippocampus and neocortex of fingolimod treated and untreated APP/PS1 mice and WT littermates.

**Suppl. Fig. 3: No increases in BDNF protein levels or TrkB signaling in fingolimod treated APP/PS1 mice**

5-6 months old male APP/PS1 mice were treated with i.p. injections of fingolimod for 1-2 months. **(A)** ELISA analysis of overall BDNF protein was performed for 6-13 animals per condition. In the hippocampus, BDNF protein levels were not significantly different between the 4 groups. BDNF protein levels in the neocortex of the same animals were in general significantly increased in APP/PS1 mice ($F(1,37) = 9.69$ $p = 0.0036$). However, there was no increase in BDNF levels in response to fingolimod treatment, neither in APP/PS1 mice nor in WT littermates (F's ≤ 0.55, p's ≥ 0.46). **(B)** Western blot analysis of TrkB tyrosine phosphorylation in the hippocampus was analyzed in 6-8 animals per condition. Fingolimod treatment resulted in slightly reduced phosphorylation of tyrosine residues Y816, Y706, and Y515 in APP/PS1 mice and WT littermates, reaching statistical significance only for Y816 ($F(1, 23) = 4.500$, $p = 0.0449$, n = 6-8 per group, two-way ANOVA). There was no effect of either genotype ($F(1,23) = 0.7829$, $p = 0.3854$) or interaction ($F(1,23) = 0.0527$, $p = 0.8205$). Western blot analysis revealed unaltered overall TrkB expression levels after fingolimod treatment in APP/PS1 mice and WT littermates (lower right). TrkB protein levels were not significantly different between WT and APP/PS1 mice (not shown).

**Suppl. Fig. 4: Dependence of CA1 pyramidal neuron spine density from distance to Aβ plaques in APP/PS1 (AD) mice.**

5-6 months old male APP/PS1 mice were treated with i.p. injections of vehicle or fingolimod for 1-2 months. Each data point represents the spine density of a dendritic segment of an individual neuron. **(A)** Plot of spine density vs. plaque distance for AD vehicle treated animals. Although there was no abrupt change in spine densities >50 µm away from plaques, spine densities were significantly smaller at <50 µm distance. The pink horizontal line represents the average spine density at distances <50 µm, and the blue horizontal line indicates the average spine density >50 µm away from the plaque border in AD vehicle mice. Classification of plaques into "near" (i.e. <50 µm) and "distant" (i.e. >50 µm away from plaque) proved useful to account best for the distance dependent spine deficit. **(B)** Plot of spine density vs. plaque distance for AD FTY720 treated animals. Spine densities were significantly increased "near" and "distant" to plaques compared to AD vehicle mice (compare Fig.2).

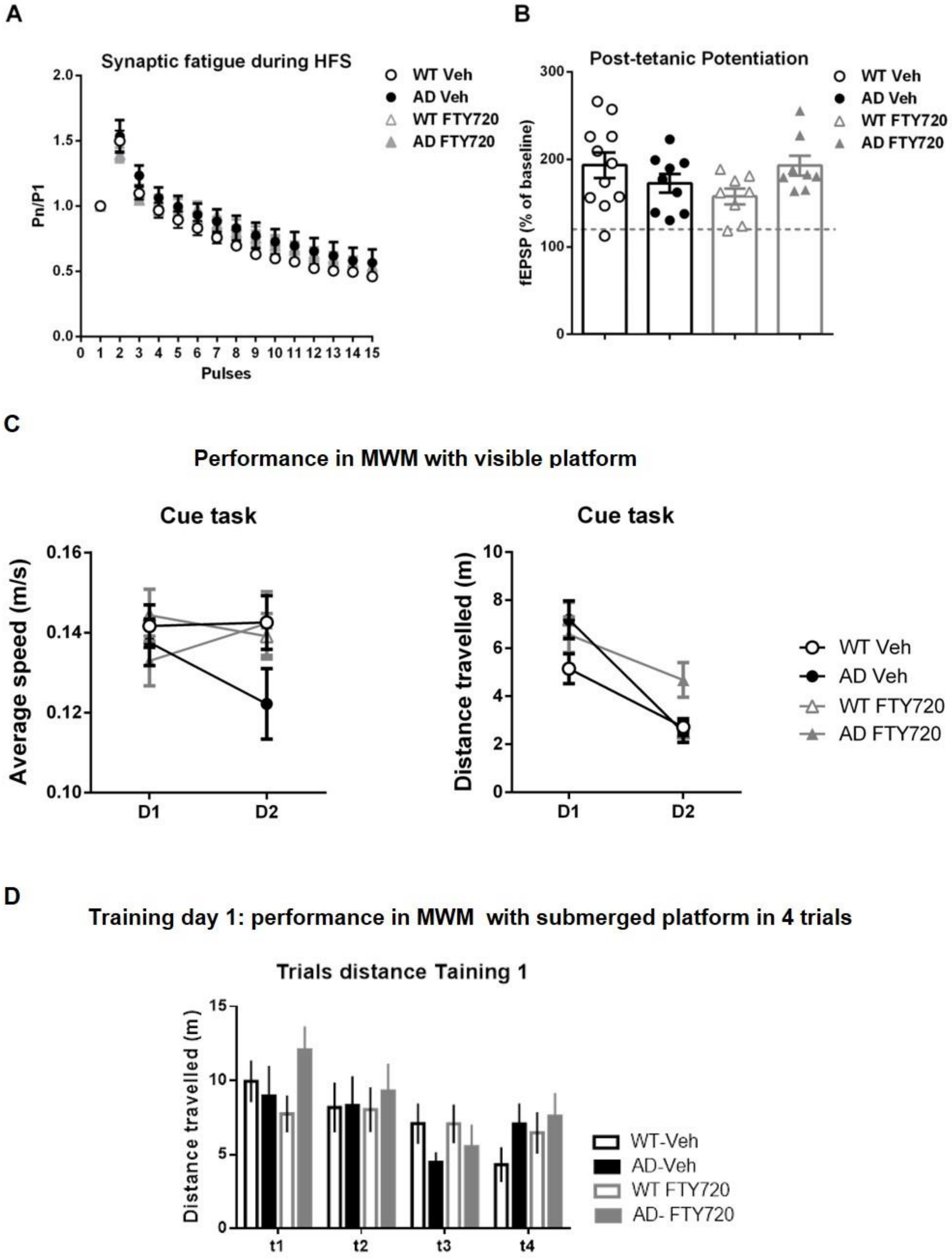
A
Synaptic fatigue during HFS
Pn/P1
Pulses
WT Veh
AD Veh
WT FTY720
AD FTY720
B
Post-tetanic Potentiation
fEPSP (% of baseline)
WT Veh
AD Veh
WT FTY720
AD FTY720
C
Performance in MWM with visible platform
Cue task
Average speed (m/s)
Cue task
Distance travelled (m)
D1
D2
WT Veh
AD Veh
WT FTY720
AD FTY720
D
Training day 1: performance in MWM with submerged platform in 4 trials
Trials distance Taining 1
Distance travelled (m)
t1
t2
t3
t4
WT-Veh
AD-Veh
WT FTY720
AD- FTY720

**Suppl. Fig. 1**

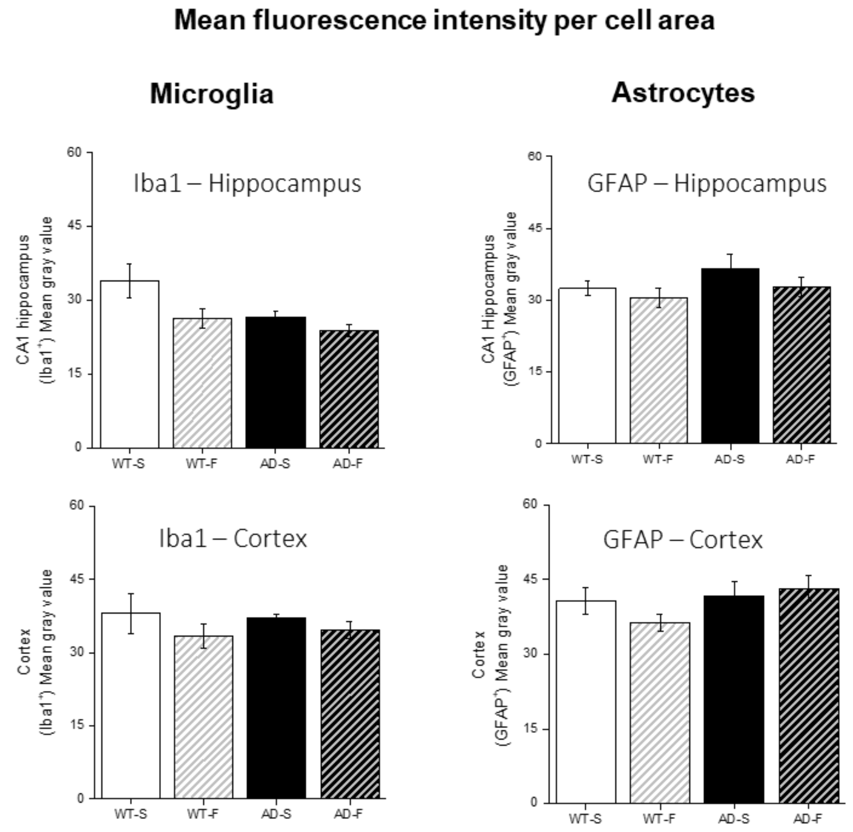


Suppl. Fig. 2

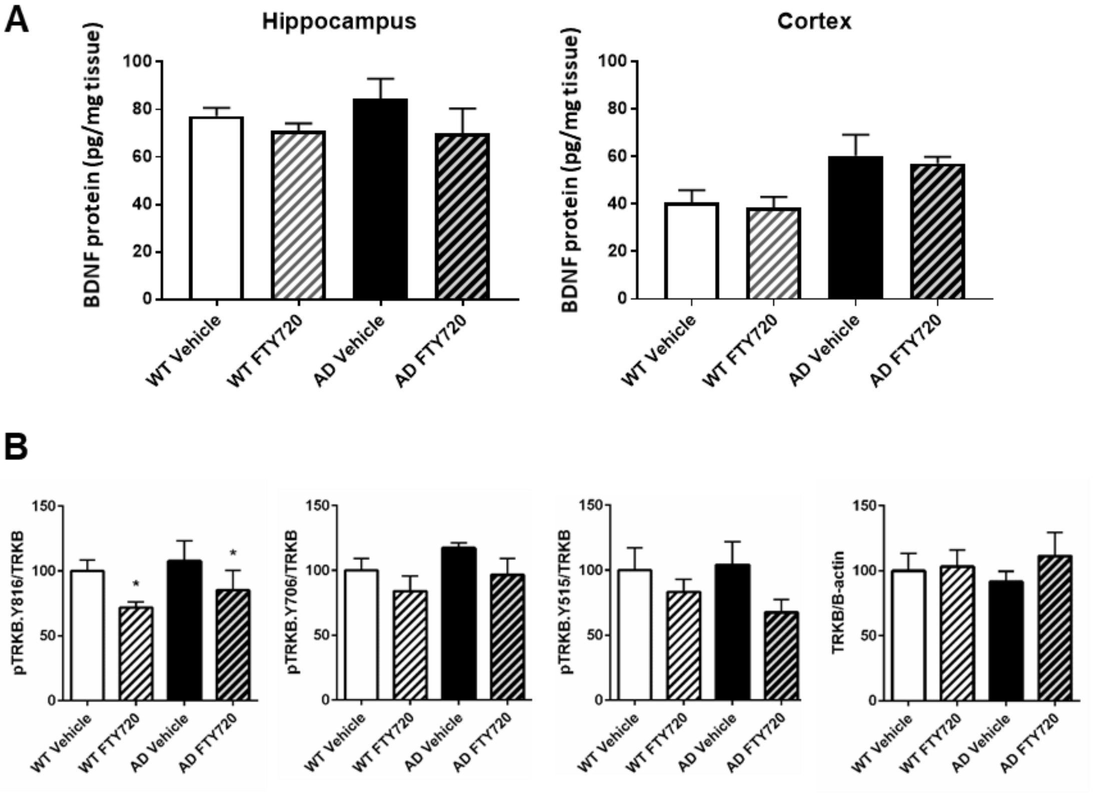


Suppl. Fig. 3

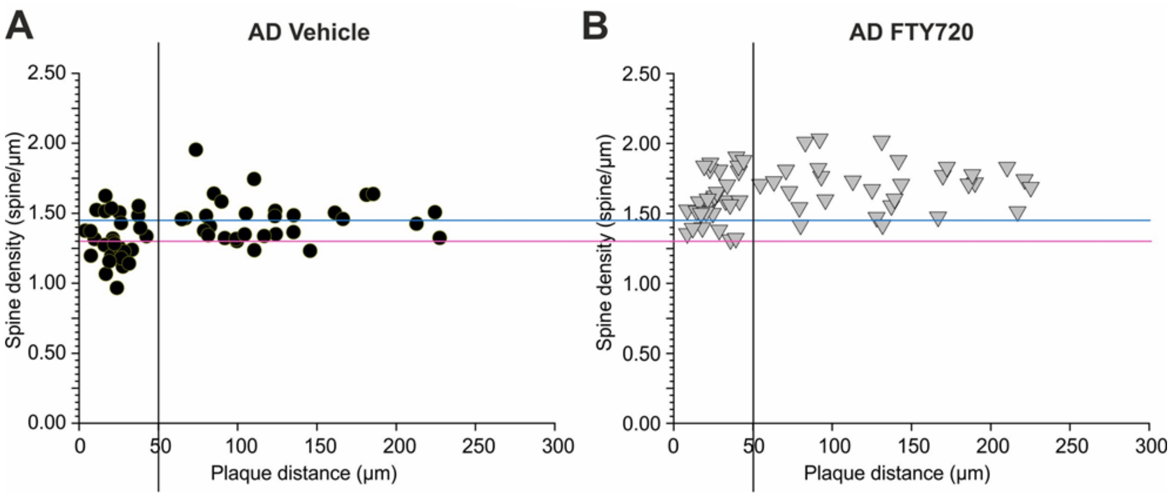
A
AD Vehicle
2.50
2.00
1.50
1.00
0.50
0.00
Spine density (spine/µm)
0
50
100
150
200
250
300
Plaque distance (µm)
B
AD FTY720
2.50
2.00
1.50
1.00
0.50
0.00
Spine density (spine/µm)
0
50
100
150
200
250
300
Plaque distance (µm)

**Suppl. Fig. 4**
